# Polyamines promote disordered protein phase separation

**DOI:** 10.1101/2024.05.10.593468

**Authors:** Matthew Percival, Christian F. Pantoja, Maria-Sol Cima-Omori, Stefan Becker, Markus Zweckstetter

**Affiliations:** German Center for Neurodegenerative Diseases (DZNE), Von-Siebold-Str. 3a, 37075 Göttingen; Max Planck Institute for Multidisciplinary Sciences, Department of NMR-based Structural Biology, Am Fassberg 11, 37077 Göttingen, Germany

**Keywords:** αSyn, Spermine, Polyamines, Metabolites, Electrostatics, Biomolecular condensates, Aggregation, Competitive Interactions

## Abstract

Membrane-less organelles are spatially heterogenous deposits of interacting macromolecules, often intrinsically disordered proteins and RNA, that form and dissolve in response to cellular stimuli. How membraneless organelles control composition while maintaining stimuli-responsiveness in an environment with competitive interactions is not well understood. Here we demonstrate that natural polyamines, which are found in all living organisms and help in many biological processes, promote protein phase separation via attractive interactions with acidic disordered domains. We show that the abundant polyamine spermine promotes phase separation of the stress-granule associated protein G3BP1 and modulates together with RNA the phase separation and amyloid formation of the Parkinson’s disease-related protein α-synuclein. Polyamine-promoted phase separation is controllable via polyamine acetylation and RNA-mediated competitive interactions. The results suggest that cellular polyamines may serve diverse roles in biomolecular condensation and the regulation of membraneless organelles.

## Introduction

Membrane-less organelles (MLOs) form via a process akin to liquid-liquid phase separation (LLPS) and are implicated in a wide range of biological processes ^1^. MLO-associated biomolecular condensation involves reversible interactions amongst macromolecules with spatially distributed stickers along the sequence ^2,3^. MLOs are spatially heterogenous, consist of many different macromolecules, and are stimuli responsive ^4,5^. The question of how MLOs control composition while maintaining stimuli-responsiveness in a cellular environment of competitive interactions is not understood.

The natural polyamines spermine, spermidine and putrescine, cellularly abundant small cationic molecules, are found in most living organisms. Their multivalent, cationic nature at physiological pH has led to the suggestion that in vivo their main state is in the bound state with nucleic acids since they are strong polyelectrolytes ^6^. Polyamines can also interact electrostatically with acidic domains of disordered proteins and, in doing so, affect protein conformations and self-assembly behaviour ^7–9^. The polyamines regulate multiple cellular processes, ranging from translation, autophagy and ion channel gating ^10–12^. This polyamine-mediated cellular regulation is critical for cellular function, and dysregulation is considered a contributing factor to cancer and neurodegenerative diseases ^13–15^.

Owing to the established connection between polyamines and nucleic acids, in addition to the plethora of intermolecular interactions between RNA and disordered proteins in MLOs ^5,16–18^, polyamines may thus have an important role in contributing to the regulation of MLOs via competitive interactions.

Here, we study the mechanistic connection between the three natural polyamines spermine, spermidine and putrescine, and RNA or disordered proteins, and their resulting phase separation behaviour. We demonstrate that polyamines promote the phase separation of the stress granule-associated protein G3BP1 and the Parkinson’ disease associated protein α-synuclein (aSyn) in a polyamine concentration and acetylation state-dependent manner. We further show that RNA sequesters polyamines from aSyn’s C-terminus, dissolves aSyn from polyamine-induced droplets and delays aSyn’s aggregation kinetics highlighting the importance of competitive interactions in the regulation of phase separation. Our study suggests an important role of natural polyamines for the regulation of membrane-less organelles.

## Results

### The cellular polyamine spermine promotes G3BP1 phase separation

G3BP1 is a central component of stress granules, MLOs which form in response to cellular stress and sequester mRNAs ^19–21^. A plethora of evidence links polyamines to stress granule regulation ^15,22–24^. However, it is challenging to determine whether polyamines are directly important for stress granule assembly or disassembly, i.e. via direct intermolecular interactions with the constituents. G3BP1 contains a large, disordered segment in the centre of its sequence which is the most acidic domain of the protein (**Figures 1A, B**). We therefore hypothesised that the strongly positively charged polyamine spermine can interact with the IDR1 region and promote G3BP1’s phase separation.

**Figure 1.**
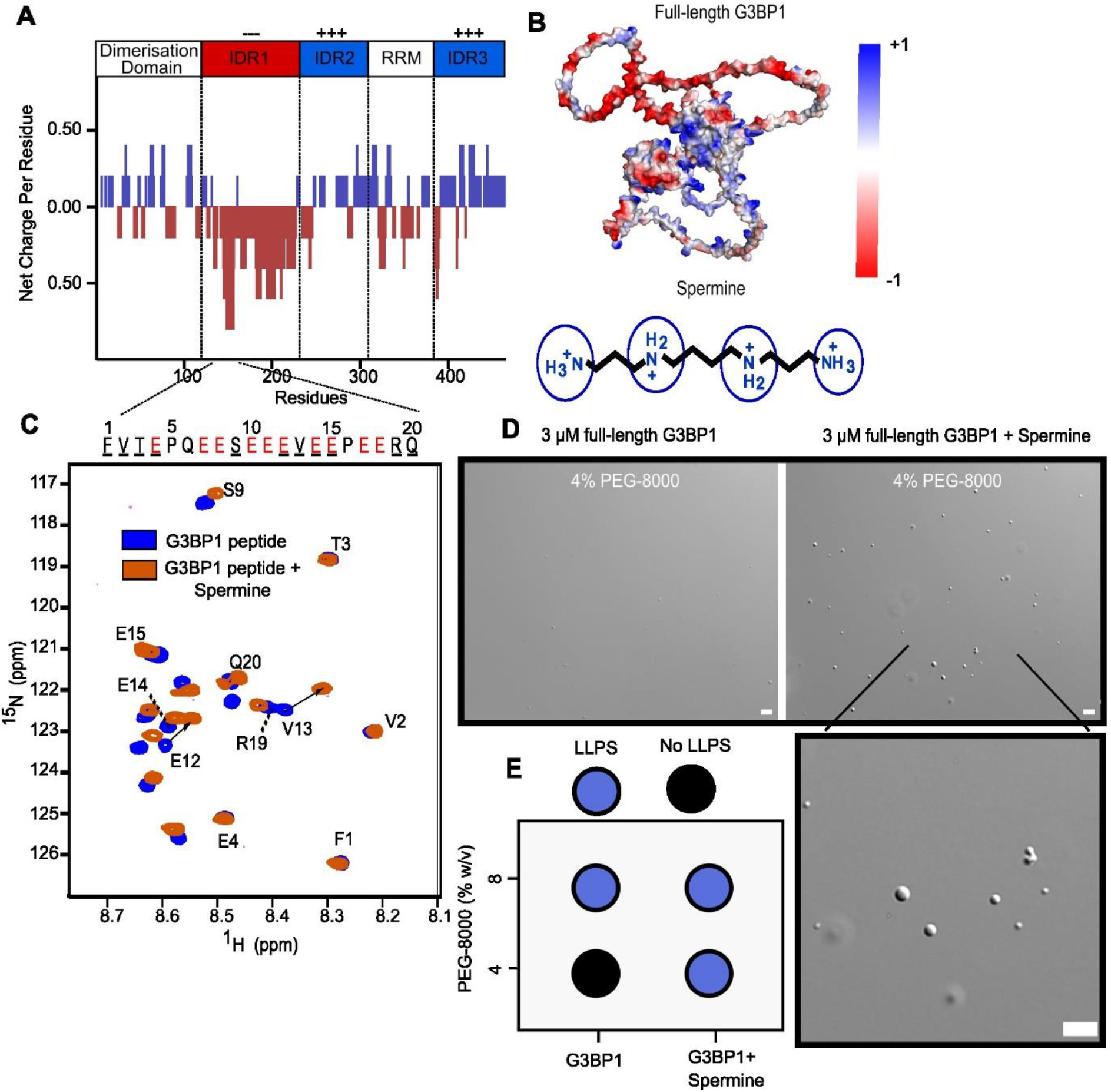
Spermine promotes G3BP1 phase separation. (A) Net charge per residue plot of full-length G3BP1. (B) Alpha Fold-2 predicted structure of G3BP1 (UniProt Q13283). Colours correspond to smoothed net charge of the protein sequence projected onto the structure. (C) ^1^H-^15^N HSQC spectrum of 500 µM G3BP1 peptide + 1250 µM spermine (orange) and G3BP1-peptide (blue) alone. G3BP1 peptide sequence is shown above spectrum, with charged residues highlighted in red and assigned residues present in the HSQC underlined in black. (D) Microscopy of 3 µM G3BP1 and 3 µM G3BP1 + spermine in 5% w/v PEG-8000. Scale bar, 5 µm. (E) Phase separation stoichiometry diagram of G3BP1 and G3BP1 + spermine at different PEG-8000 concentrations.

To determine whether spermine can directly interact with the IDR1 region of G3BP1, we synthesized a 20 amino acid peptide from G3BP1’s IDR1 (G3BP1-pep). The NMR signals of G3BP1-pep were sequence specifically assigned using a combination of two-dimensional (2D) TOCSY and NOESY experiments. Subsequently, spermine was mixed with the peptide and a 2D ^1^H-^15^N HSQC recorded **(Figure 1C)**. Comparison of the ^1^H-^15^N HSQC without spermine revealed many spermine-induced chemical shift perturbations. Sequence-specific analysis assigned the strongest perturbations to the acidic C-terminus of G3BP1-pep, and conversely very few to the N-terminal residues **(Figure S1A)**. Residues E12 and V13 (corresponding to G3BP1 residues E152 and V153) displayed the largest chemical shift perturbations, likely highlighting their positions at the centre of the spermine interaction site.

Next, we performed full-length G3BP1 phase separation experiments without and with spermine. Firstly, we incubated 5 µM G3BP1 in 8 % w/v PEG-8000 solution. Phase contrast microscopy confirmed the presence of droplets **(Figure S1B)**. Similar sized droplets were observed in the presence of spermine **(Figure S1)**. We then decreased the G3BP1 concentration to 3 µM and the PEG-8000 concentration to 5 % (**Figure 1D)**. In these less phase separation-prone conditions, little if any phase separation was observed without spermine **(Figure 1D, left panel)**. However, the addition of spermine generated more droplets of larger size **(Figure 1D, right panel)**. Turbidity measurements reinforced these findings (**Figure S1C**). The data demonstrate that the cellular polyamine spermine promotes the phase separation of the stress granule-scaffold protein G3BP1.

### aSyn and spermine form clusters in solution

Next, we investigated the interaction of the naturally occurring polyamine spermine with the intrinsically disordered 140-residue protein aSyn. NMR spectroscopy revealed an interaction site in the acidic C-terminus of aSyn (**Figure 2A**). In agreement with previous reports, the interaction site is centred around the most acidic region of the C-terminus comprising residues D119-E136 and the affinity is in the micromolar range ^9^ **(Figure 2A, inset)**.

**Figure 2.**
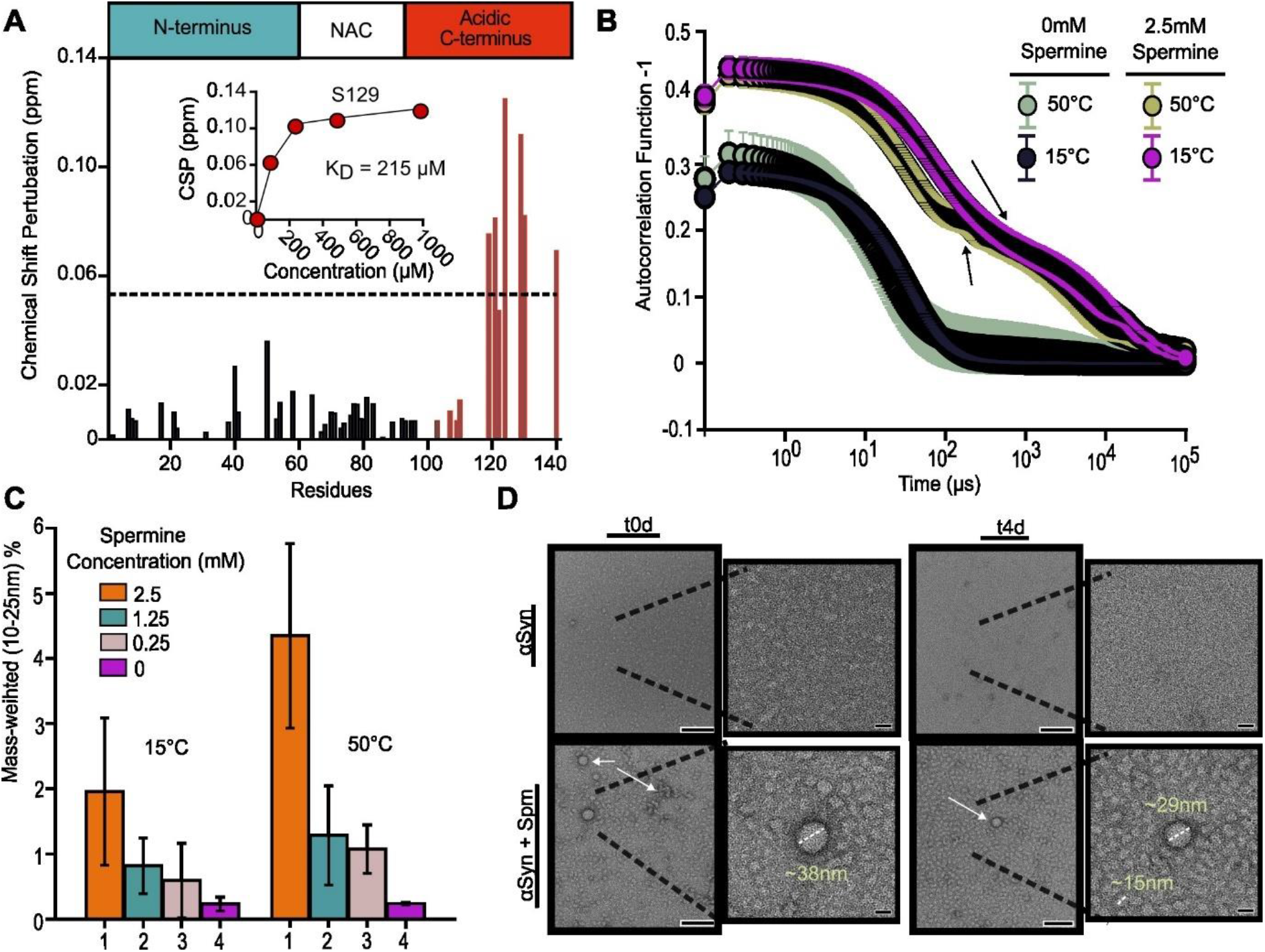
Spermine stabilises aSyn clusters. **(A)** Chemical shift perturbation of 50 µM aSyn plus 1 mM spermine. Black dotted line corresponds to 2 standard deviations of chemical shift perturbations along the entire sequence. Inset shows chemical shift perturbation of S129 with increasing spermine concentration. K^D^ value is the best fit of non-linear regression with 95% confidence intervals 116 to 390 µM. **(B)** Autocorrelation function of 250 µM aSyn and 250 µM aSyn + 2.5mM spermine at 15°C and 50°C. Samples were incubated at relevant temperature for 5 minutes before the start of the measurement. Black arrows point toward wiggle and oscillations in the slow decaying part of the curve. **(C)** Mass-weighted intensity increase of 250 µM aSyn and aSyn with various molar ratios with spermine between the radii 10 and 25 nm at 15°C and 50°C. Error bars represent standard deviation. **(D)** Transmission Electron Microscopy (TEM) micrographs of 250 µM aSyn (top row) and aSyn + 2500 µM Spermine (bottom row) at room temperature. First two columns correspond to measurements at t0d whereas the latter two columns correspond to measurements of the sample sample but at t4d. Scale bar of original images 100nm, scale bar of zoomed images 20nm.

Phase separation between aSyn and spermine was tested using a temperature-ramp dynamic light scattering (DLS) experiment. At 250 µM aSyn and 2500 µM spermine, bimodal decays and ‘wiggles’ (**Figure 2B**, arrows) present in the autocorrelation function at 15 °C and 50 °C indicated the presence of inhomogeneities likely representing high molecular weight structures **(Figure 2B)**. In contrast, without aSyn present the decay was monomodal and representative of small molecule tumbling, such as spermine **(Figure S2A)**. The mass-weighted median radius of aSyn and spermine was 12.1 nm compared to 5.1 nm without spermine present **(Figure S2B)**. These averaged radii are much larger than monomeric aSyn and reflect the presence of high molecular weight species coexisting with monomeric protein. Size analysis indicated that the high molecular weight species are in the range of 10-25 nm in diameter **(Figure 2C)**. The mass fraction of high molecular weight species with both aSyn and spermine present was higher at 50 °C compared to 15 °C but was not statistically significant and showed a strong dependence on aSyn: spermine molar ratio **(Figure 2C)**. This stoichiometry-dependency is to be expected if the high molecular weight species that are formed are correlated to the heterotypic interaction of spermine with the C-terminus of aSyn.

To confirm the presence of high molecular weight structures, samples were imaged with transmission electron microscopy. Electron microscopy revealed oligomers of various shapes **(Figure 2D)**. At room temperature and 50 °C, oligomers were present in both aSyn and aSyn/spermine samples **(Figure 2D and Figure S2D)**. However, the average size and extent of the oligomers grew substantially in the presence of spermine **(Figure 2D**, bottom row). In contrast, without aSyn the micrographs were sparsely populated **(Figure S2C)**. The micrographs containing spermine and aSyn additionally displayed the presence of larger spherical objects (diameter of ∼32 nm) (**Figure 2D, arrows, inset)**. However, most of the objects were not spherical and appeared to have more complex shapes **(Figure 2D)**. Interested in the time-dependent evolution of the clusters, we measured the same samples after an incubation of four days at room temperature. In the case of aSyn and spermine, the size homogeneity of the oligomeric particles apparently increased during the incubation period **(Figure 2D, lower right)**. In contrast, the micrograph with only aSyn was more sparse after incubation for four days when compared to the start of incubation **(Figure 2D, upper right** compared to **Figure 2D, upper left)**, suggesting that the oligomeric particles are unstable without spermine.

Previous reports demonstrated that aSyn can form nanoclusters in high ionic strength and crowding conditions that evolve over-time into a macroscopic phase separated state ^25^. We did not observe time-evolution into a macroscopic phase separated state; in contrast, the clusters were more stable with spermine than without. This increased stability may be explained by an excess charge present in the clusters that prevents growth into a macroscopic state. We propose that a combination of increased effective attraction between αSyn monomers mediated by spermine increases the extent of the clusters, and the excess charge prevents further growth thus stabilizing a microscopic state. This type of mixed potential is often seen in systems generally with mixed attractive and repulsive interactions ^26^.

### Spermine promotes phase separation of aSyn

To investigate whether aSyn and spermine can also form macroscopically phase separated states, we added the macromolecular crowder PEG-8000 to the solution. At the concentrations tested, neither aSyn alone, nor aSyn with only spermine, nor spermine with PEG underwent phase separation ^27^ (**Figure S3A)**. Similarly, aSyn with only PEG did not phase separate, in agreement with previous reports **(Figure S3A)** ^27^. However, in the presence of spermine and 20 % PEG turbidity measurements revealed sigmoidal growth profiles, with maxima occurring at approximately a 1:5 to 1:10 molar ratio of aSyn to spermine (**Figure 3A**). Microscopy experiments revealed the presence of aSyn-rich droplets in a spermine concentration-dependent manner highlighting the aSyn: spermine stoichiometry is important for the phase separation (**Figure 3B**). Additionally, phase separation between aSyn and spermine was observed without fluorescently labelled aSyn **(Figure S3B)**. aSyn-rich droplets were detected at 20 µM but not 10 µM, and their total area and frequency increased at 50 µM **(Figure S3 C**,**D)**. Therefore, the lowest concentration for phase separation of aSyn in the presence of spermine and 20 % PEG is likely between 10 and 20 µM. At the lower PEG concentration of 10 % w/v, the lowest concentration increased to approximately 200 µM **(Figure S3E)**. At 5 % PEG, no phase separation was observed within the concentration ranges used (**Figure S3E)**.

**Figure 3.**
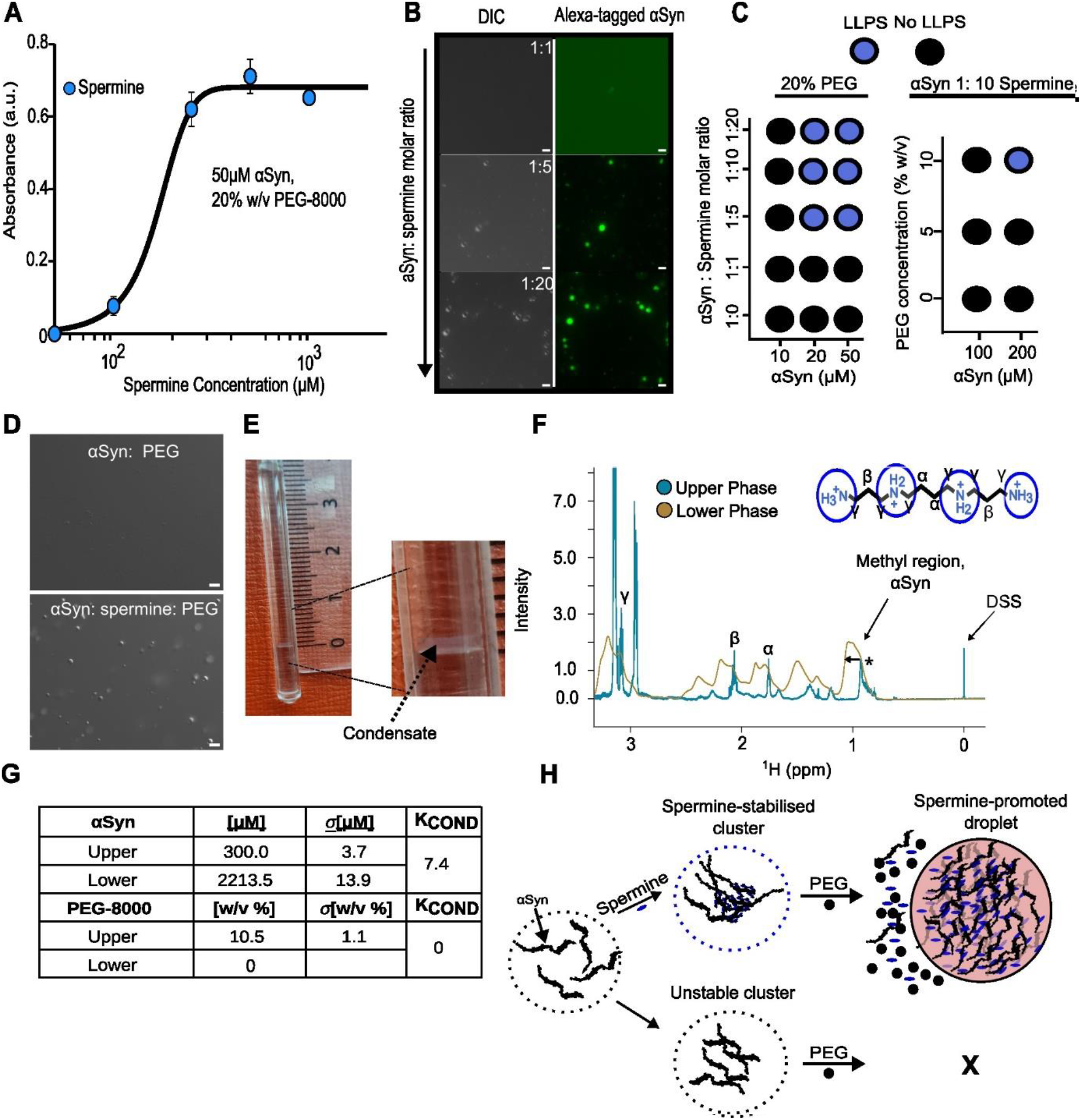
αSyn is enriched and PEG depleted in spermine-dependent condensate. **(A)** Turbidimetric plot using a wavelength of 600nm. aSyn was held constant at a 50 µM concentration and spermine added at various concentrations. **(B)** DIC (left column) and fluorescence (right column) microscopy of aSyn with various molar ratios of spermine. Scale bar, 5 µm. **(C)** Stoichiometry-dependence of phase separation. **(D)** DIC micrographs of aSyn with only PEG and aSyn with PEG and spermine (lower) **(E)** Picture of aSyn condensate inside Shigemi NMR tube **(F)** 1H NMR spectrum of condensate (lower) and dilute phase (upper) **(G)** Concentration quantification (H) Schematic of process. Scale bars 5 µm

To determine the composition of the phase separated system we used spatially-resolved NMR ^28^. This high-resolution method allows us to discriminate signals in a position-dependent manner between those in the concentrated phase (condensate, lower phase) and the dilute phase (upper) phase. In conditions of 400 µM aSyn, 4mM spermine and 10% w/v PEG-8000 we observed an abundance of µm-sized droplets using phase contrast microscopy **(Figure 3D, bottom)**. For the NMR experiment we increased the concentration of aSyn to 600 µM and maintained the PEG concentration at 10% w/v and the aSyn: spermine molar ratio at 1: 10. Phase separation of only aSyn was not observed even at 800 µM concentration **(Figure 3D, top)**. After centrifugation of the aSyn, spermine and PEG solution a 0.7 mm tall turbid region was formed at the bottom of the Shigemi NMR tube indicating the formation of an aSyn-rich condensate **(Figure 3E)**.

After equilibration of the sample at 15^0^C we performed a 1D echo imaging experiment to map the signal intensity of the water signal along the z-axis **(Figure S4A)**. For aSyn which forms a single homogenous phase in these conditions the profile is analogous to a bell or hat representing a uniformly mixed solution **(Figure S4A)**. In the case of aSyn: spermine and PEG an intensity dropped occurred close to the bottom of the Shigemi NMR tube **(Figure S4A)**. The intensity drop suggests the presence of a second phase with less water than the first phase. We then recorded 1D spatially resolved NMR spectra in the condensate (lower) and upper phases of the sample **(Figure 3F)**. The signals from the condensate were severely broadened which are a consequence of magnetic susceptibility differences between both phases **(Figure 3F, asterisks, arrow)**. The signals from the methyl region (0.6 – 1.1ppm) which contains signals only from aSyn, as well as the methylene region which contains signals from the alpha and beta protons of spermine **(Figure 3F)**, appear more intense in the condensate, whereas that of DSS (sodium trimethylsilylpropanesulfonate) – used as a chemical shift calibration reference is severely broadened almost beyond detection **(Figure 3F)**.

To determine how the composition changes from the condensate to the upper phase we recorded spectra at different z-positions (offset frequencies) of the NMR tube in the sample. The analysis determined that aSyn’s signal intensity plateaus in condensate before decreasing at higher positions which have larger contributions from the upper phase **(Figure S4B)**. Therefore, aSyn is present in the homogenous phase of the condensate. After sufficient increments in the position of the 1.5 mm excitation slice in the z-direction both the condensate and the upper phases are excited, and this is the regime where the PEG-8000 signal appears **(Figure S4C)**. In contrast to the aSyn signals which have reached a plateau, the PEG-8000 -CHsignal was substantially enhanced **(Figure S4C)**. Therefore, since the PEG-8000 signal intensity positively correlates with extent of upper phase excitation we concluded that PEG-8000 is excluded from the condensate, consistent with previous reports ^25^.

To quantify the concentrations of the solutes in the two phases we recorded reference spectra of aSyn, PEG and spermine following the protocol presented previously ^28^ **(Figure S4 D, E)**. The analysis determined that the concentration of aSyn in the dilute and condensed phases was 300.0 ± 3.7 µM and 2212.5 ± 13.9 µM, respectively **(Figure 3G)**. This yields a partition coefficient for aSyn – defined as the quotient of concentrations in the condensate and dilute phase – of 7.4. In contrast, using the resolved -CH-signal of PEG we determined its concentration in the dilute phase to be 10.5 ± 1.1% w/v (**Figure 3G**). Quantifying spermine’s concentration had lower accuracy since its detectable ^1^H signals originate from its methylene -CH2-groups which are also present in the side chains of some amino acids, for example lysine which is abundant in aSyn’s N-terminus. Despite this, we were able to determine the upper phase concentration of spermine to be 5.4 ± 0.8 mM. Since this is lower than the initial mixing conditions of 6 mM it suggests that spermine is abundant in both phases and is probably slightly enriched in the αSyn-enriched condensate.

Therefore, PEG-8000 is excluded from the condensate, whereas aSyn is majorly and spermine minorly enriched. These calculations are consistent with a segregative phase separation process between aSyn: spermine and PEG-8000 (**Figure 3H**).

### Spermine-promoted aSyn phase separation accelerates aSyn aggregation

It is known that the polyamines and especially spermine can accelerate the aggregation kinetics of aSyn ^29^. How the aggregation kinetics in the phase separated solution compare is less clear. In phase separated conditions of 50 µM aSyn, 500 µM spermine and 20 % w/v PEG-8000, the lag phase decreased substantially compared to aSyn and PEG mixtures **(Figure 4A)**. The lag phase of aSyn fibrillization decreased to ∼10 hours with both spermine and PEG present. In contrast, with only PEG present the lag phase was ∼40 hours, i.e. ∼4 times longer (**Figure 4A**). Under the same aggregation conditions but with only aSyn and spermine or with only aSyn, no thioflavin-T (ThT) positive aggregation of the protein was detected during the incubation period (**Figure 4A**).

**Figure 4.**
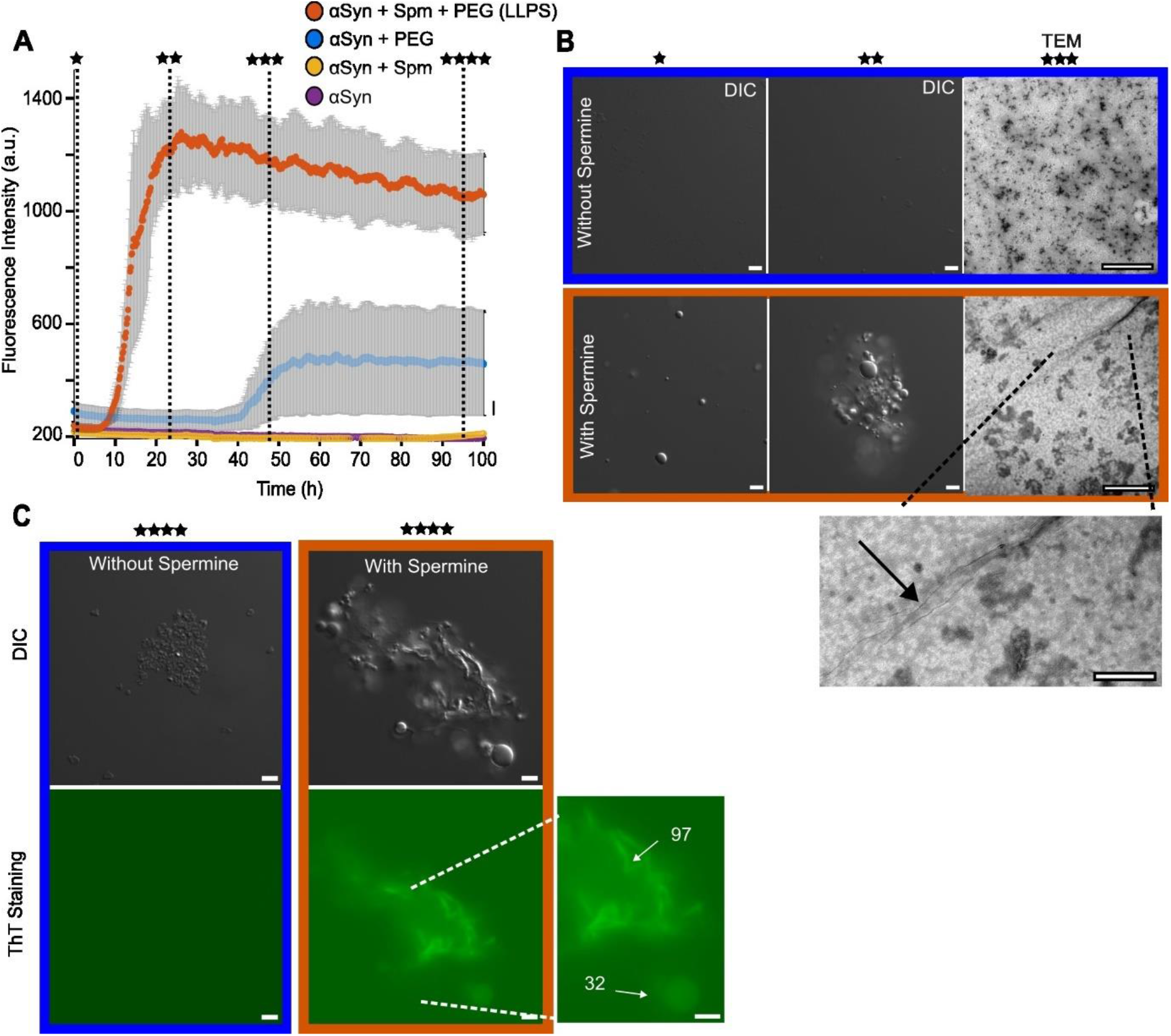
Phase separation conditions accelerate αSyn’s aggregation kinetics. **(A)** ThT aggregation assay of aSyn (50 µM) with either PEG, Spermine or PEG + Spermine, the latter being phase separation conditions. **(B)** DIC (scale 5 µm) and TEM micrographs (100 nm) of 50 µM aSyn with or without 500 µM spermine recorded at different time points. **(C)** DIC and fluorescence micrographs of aSyn and aSyn + spermine at t 4d. Fluorescent signal originates from ThT which was spiked into the aSyn and spermine samples after 4 days of incubation.

To confirm the presence of aggregates, the samples were imaged with transmission electron microscopy at different time points of the aggregation process. At the early stages, phase separation but no aggregation of the aSyn/sperm-ine/PEG sample was evident, while without spermine the sample was clear **(Figure 4B)**. After ∼24h when the ThT intensity of the phase separated sample reached saturation, phase contrast microscopy revealed both aggregates and droplets, while without spermine the sample was again clear **(Figure 4B, second column)**. At 48h of incubation, transmission electron microscopy revealed aggregates in both cases, but under the conditions of phase separation long and fibrillar species were present **(Figure 4B, third column)**. To confirm that these species were beta-sheet rich species, a low concentration of ThT was spiked into the samples. Fluorescence microscopy revealed ThT staining for both the aSyn-rich droplets and the fibrillar aggregates, albeit approximately three times stronger in the fibrillar species **(Figure 4B, second column and inset)**. In contrast to the phase separated sample, aSyn and PEG alone showed no ThT staining **(Figure 4B, first column)**. The analysis demonstrates that spermine-assisted phase separation of aSyn into proteinrich droplets accelerates its aggregation kinetics into fibrillar species that coexist with aged droplets.

### Polyamine type and ionic strength control amount of αSyn phase separation

Inside cells, spermine can be back-converted to spermidine and putrescine, two shorter but biologically abundant polyamines ^30^ (**Figure 5A**). Moreover, mono- and di-acetylated forms of spermine often form en-route to its backconversion to spermidine and putrescine (**Figure 5A**) ^12^. Acetylation of polyamines has been suggested to be an important mechanism of polyamine control since aberrant acetyltransferase behaviour results in multiple cellular problems ^30,31^. The major physicochemical consequence of back-conversion and acetylation is that it reduces the net charge of the polyamine and increases its hydrophobicity, thus weakening the electrostatic potential for interaction with oppositely charged macromolecules.

**Figure 5.**
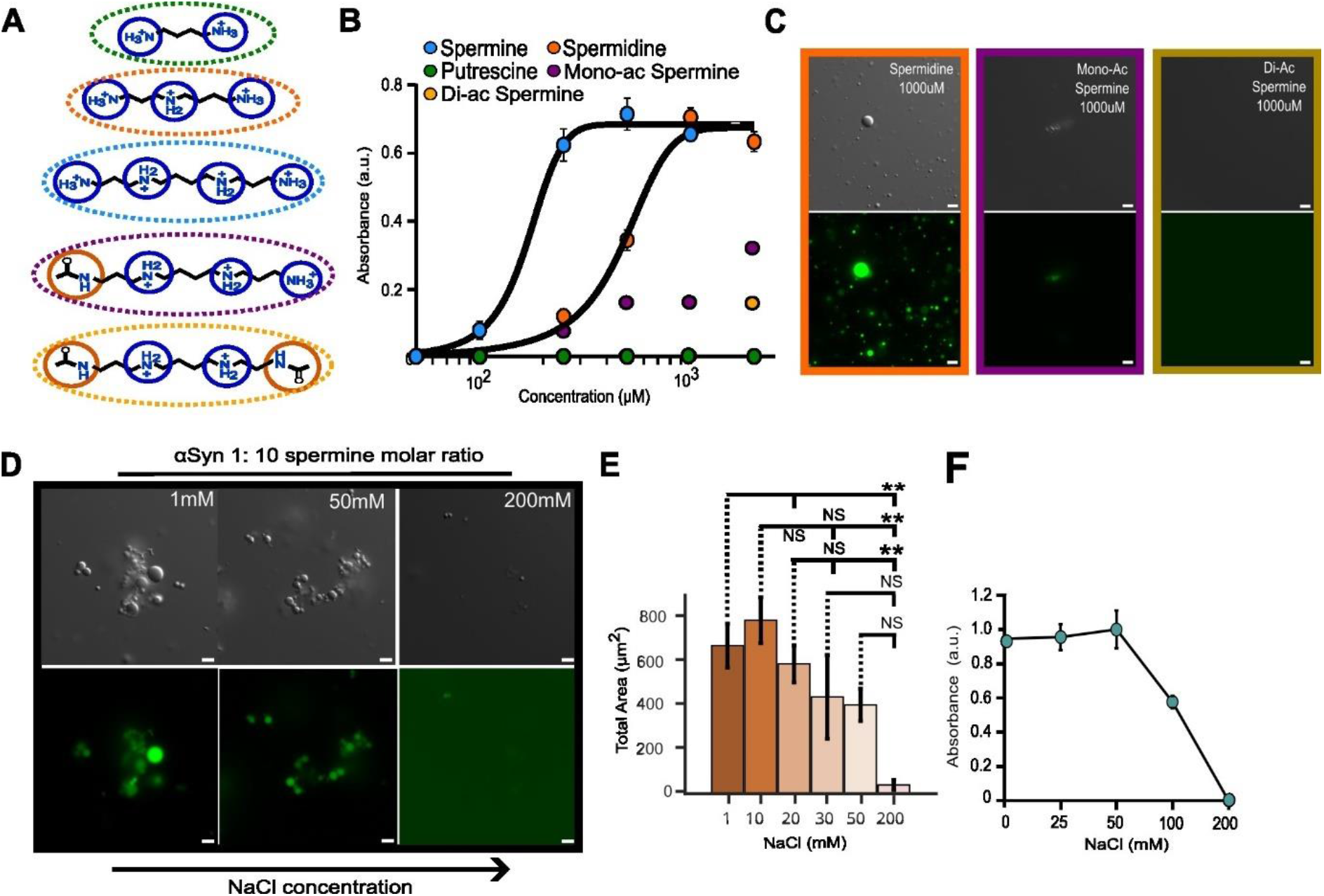
Electrostatics control αSyn: polyamine phase separation. **(A)** Chemical structures of different polyamines (top to bottom: putrescine, spermidine, spermine, mono-acetyl spermine, di-acetyl spermine). **(B)** Turbidity measurements (600 nm) of polyamineand concentration-dependent aSyn phase separation. **(C)** DIC and fluorescence micrographs of 50 µM aSyn with either 1 mM spermidine (left), 1 mM mono-acetyl spermine (second column) and 1 mM di-acetyl spermine (third column). **(D)** DIC and fluorescence micrographs of 100 µM aSyn + 1 mM spermine at various NaCl concentrations. **(E)** Total fluorescent areas of aSyn, spermine and NaCl micrographs. Error bars represent standard deviation of n=2 total area calculations. Statistical significance tests were conducted using one-way ANOVA. **(F)** Turbidity (600 nm) of aSyn + spermine at various NaCl concentrations. All scale bars 5 µm.

While the interaction site at the acidic C-terminus of aSyn remains identical for each of the polyamines, the interaction strength decreased in the order spermidine > mono-acetyl spermine > putrescine > di-acetyl spermine which correlates with the decreasing net charge of the polyamines (**Figure S5A**). In comparison to spermine, higher concentrations of each of the polyamines were necessary to phase separate aSyn **(Figure 5B)**. At a 1: 20 molar ratio only spermidine produced a significant volume of the phase separated state **(Figure 5C)**. Spermidine, bearing approximately 2.7 protonated groups at neutral pH, led to aSyn-rich droplets at a 1:5 molar ratio and higher **(Figure S5B)**. At a molar ratio of 1:1, however, no phase separation was observed **(Figure S5B)** Putrescine, bearing slightly less than two protonated groups at neutral pH, required a 100-fold molar excess to phase separate aSyn **(Figure S5B)**. This reflects the much weaker binding affinity of putrescine to aSyn.

Acetylation of spermine also led to substantially weaker phase separation with aSyn **(Figure 5C)**. For di-acetyl spermine, no phase separation was observed at any concentration **(Figure 5C)**. Partly this is due to the much weaker binding affinity of di-acetyl spermine to aSyn compared to the other polyamines, but also it reflects the poor solubility of the compound in water as higher concentrations were unreachable without precipitation. The ability to promote aSyn phase separation was thus ordered spermine > spermidine > mono-acetyl spermine > putrescine > di-acetyl spermidine. The metabolic conversion of spermine and physiologically essential acetylation of spermine’s primary amines thus reduces the interaction strength and phase separation with the paradigmatic disordered protein aSyn, highlighting the importance of electrostatics.

If electrostatics are dominating the heterotypic phase separation between αSyn and the polyamines, there should be a reduction in phase separation volume with increasing ionic strength. At 100 µM aSyn and 1000 µM spermine, addition of up to 50 mM NaCl did not significantly affect total droplet area **(Figure 5D,C)**. However, at 200 mM NaCl turbidity was absent and minimal aSyn-rich objects were observed with microscopy (**Figure 5D-F**). The sensitivity of polyamine-promoted phase separation to salt concentration is analogous to the influence of ionic strength on complex coacervation whereby strong polyelectrolytes, which interact electrostatically, phase separate ^32^. This is despite the phase separation process in our system undoubtedly involving a combination of heterotypic, homotypic, and crowding-induced interactions that involve ion-pairs, hydrogen bonds, Van der Waal’s interactions, and hydration effects that are non-uniform along the sequence. While therefore it may be too simplistic to label this process as complex coacervation, it nevertheless highlights the importance of the heterotypic electrostatics for the phase separation in this system.

Our results add to previous reports on aSyn phase separation that demonstrated high ionic strength can lead to phase separation of the protein ^25,36^. Our results differ because of the presence of spermine to promote a heterotypic phase separation which circumvents the necessity of high ionic strength to promote the homotypic phase separation. This also avoids the perturbation of bulk hydration expected at high ionic strength. Therefore, direct and predominantly electrostatic interactions between the polyamines and the acidic C-terminus of aSyn drive the condensation process, and changing the strength of this interaction affects the amount of phase separation. This is consistent with previous reports on the distinction between heterotypic and homotypic phase separation of charged biopolymers ^34^.

### RNA sequesters polyamines from the aSyn C-terminus

RNA is abundant in MLOs where it forms a variety of intermolecular interactions with stickers of disordered proteins ^17,20,35^. Since polyamines interact strongly with RNA and can promote its phase separation ^36^, we investigated the interaction equilibria of polyU RNA, spermine and aSyn and its relation to phase separation.

To study the interaction in more detail, we followed the ^1^H NMR signal of the alpha proton of spermine at 15 °C to prevent phase separation **(Figure S6 A**,**B)**. Titrating polyU RNA into a 250 µM spermine solution led, at low RNA concentrations, to lower chemical shift values of spermine’s methylene protons indicative of an interaction **(Figure S6C, left panel)**. At higher concentrations a combination of large chemical shift changes alongside broadening of spermine’s signals occurred **(Figure S6C, D, left panel)**. The broadening of spermine’s chemical shifts indicates a strong interaction between RNA and spermine. The origin of the broadening can either be from chemical exchange effects or an increased transverse relaxation rate of spermine in the bound state with RNA.

Since spermine interacts with both RNA and aSyn, we next investigated a ternary system of spermine, aSyn and RNA **(Figure S6A, right)**. If RNA has a much higher binding affinity for spermine than aSyn we expected a similar effect to spermine’s chemical shifts with increasing RNA concentration; moreover, we expected the spermine-induced chemical shift perturbations of aSyn to return to their reference values reflecting a decreased population of sperminebound aSyn.

In the presence of aSyn, spermine’s methylene proton signals show a similar pattern to what was observed with only RNA in solution, namely lower field chemical shift changes followed by signal broadening at higher RNA concentrations **(Figure S6C, right)**. This analysis suggests that the interaction between polyU RNA and spermine is qualitatively similar with or without aSyn present. We also recorded HSQC spectra of aSyn in the same conditions to determine if RNA is decreasing the population of the aSyn: spermine bound state **(Figure 6A)**. The analysis determined that aSyn’s chemical shifts in the presence of spermine return to their reference values with increasing RNA concentration **(Figure 6B, C)**. The chemical shifts do not return exactly to the reference chemical shifts of aSyn indicating that there is still some spermine correlated to the C-terminus at high polyU RNA concentration. There was no evidence of broadening of aSyn’s chemical shifts, nor of chemical shift perturbations in the lysine-rich N-terminus of the protein. These observations suggest that RNA – as a consequence of its much higher binding affinity for spermine than aSyn ^6^ – is sequestering spermine from the C-terminus of aSyn.

**Figure 6.**
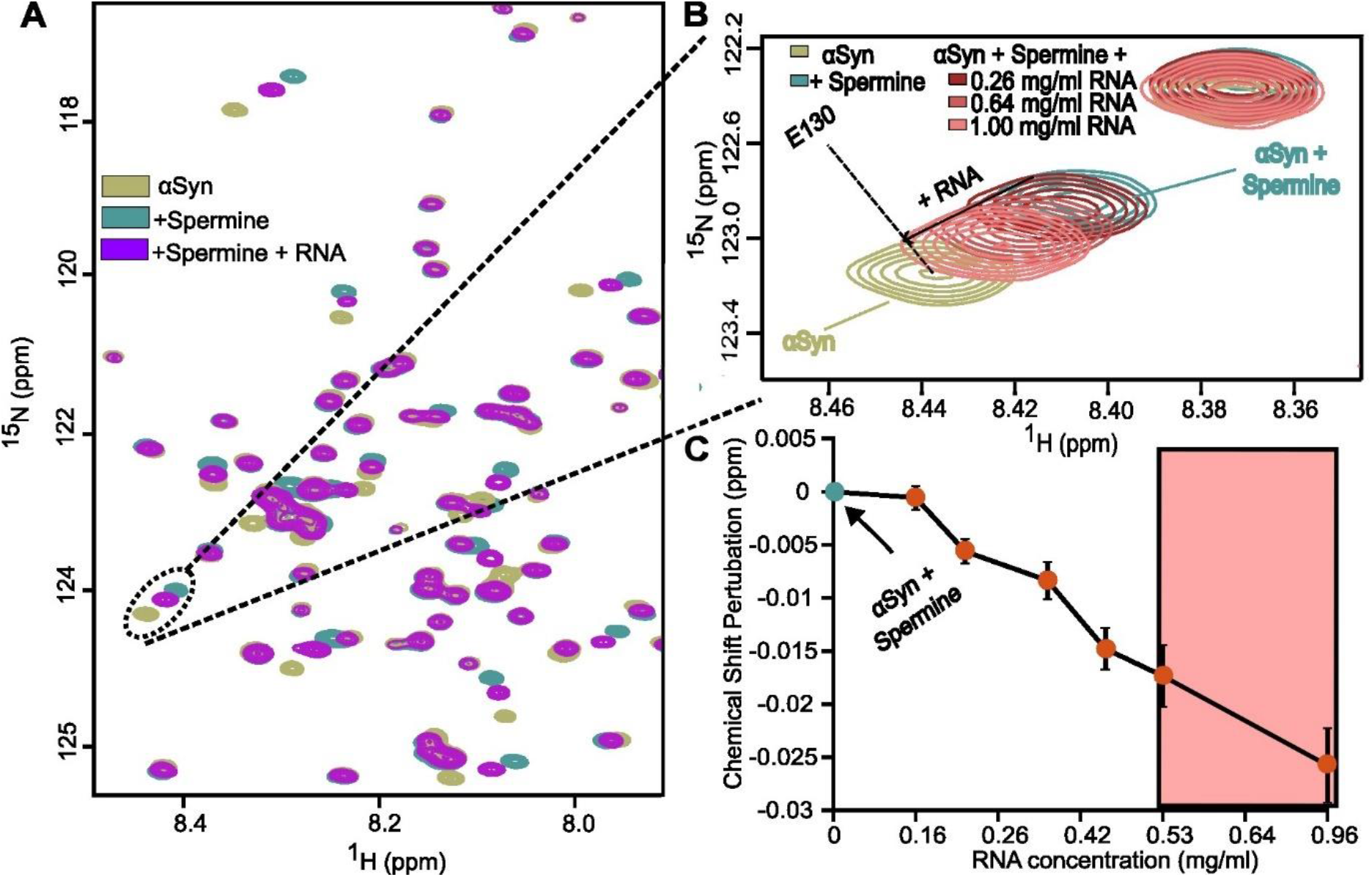
RNA sequesters spermine from aSyn’s C-terminus. **(A)** ^1^H-^15^N HSQC spectra of 50 µM ^15^N-labelled aSyn (olive) in the presence of constant 500 µM spermine (aqua) and increasing concentrations of polyU RNA (purple). **(B)** Inset of HSQC showing aSyn residue E130 undergoing chemical shift perturbations in the presence of increasing polyU RNA in solution. **(C)** Chemical shift perturbation of aSyn + spermine with increasing polyU RNA concentrations. Data points represent the average CSP of four aSyn residues, S129, E130, E135 and A140 at each RNA concentration. Error bars represent standard deviation of the chemical shift perturbations.

### RNA resolubilises αSyn

In microscopy-based phase separation experiments, we observed that with increasing RNA concentration the intensity of aSyn’s fluorescence decreased substantially **(Figure 7A)**. At the lowest RNA concentration used, aSyn remained strongly concentrated in the spermine-dependent droplets **(Figure 7A, top row)**. At higher polyU RNA concentrations, however, the fluorescence intensity decreased **(Figure 7B)** and a small number of shell-like droplets became present **(Figure S7A)**. At these high RNA concentrations, aSyn appeared to partition to the interface of some droplets **(Figure 7A, bottom row)**. The results indicate that polyamine-promoted phase separation of aSyn is readily controllable through competitive interactions involving RNA. They highlight the importance of the heterotypic electrostatics between aSyn’s C-terminus and spermine and suggest a mechanism of resolubilisation of charged biopolymers via competitive interactions.

**Figure 7.**
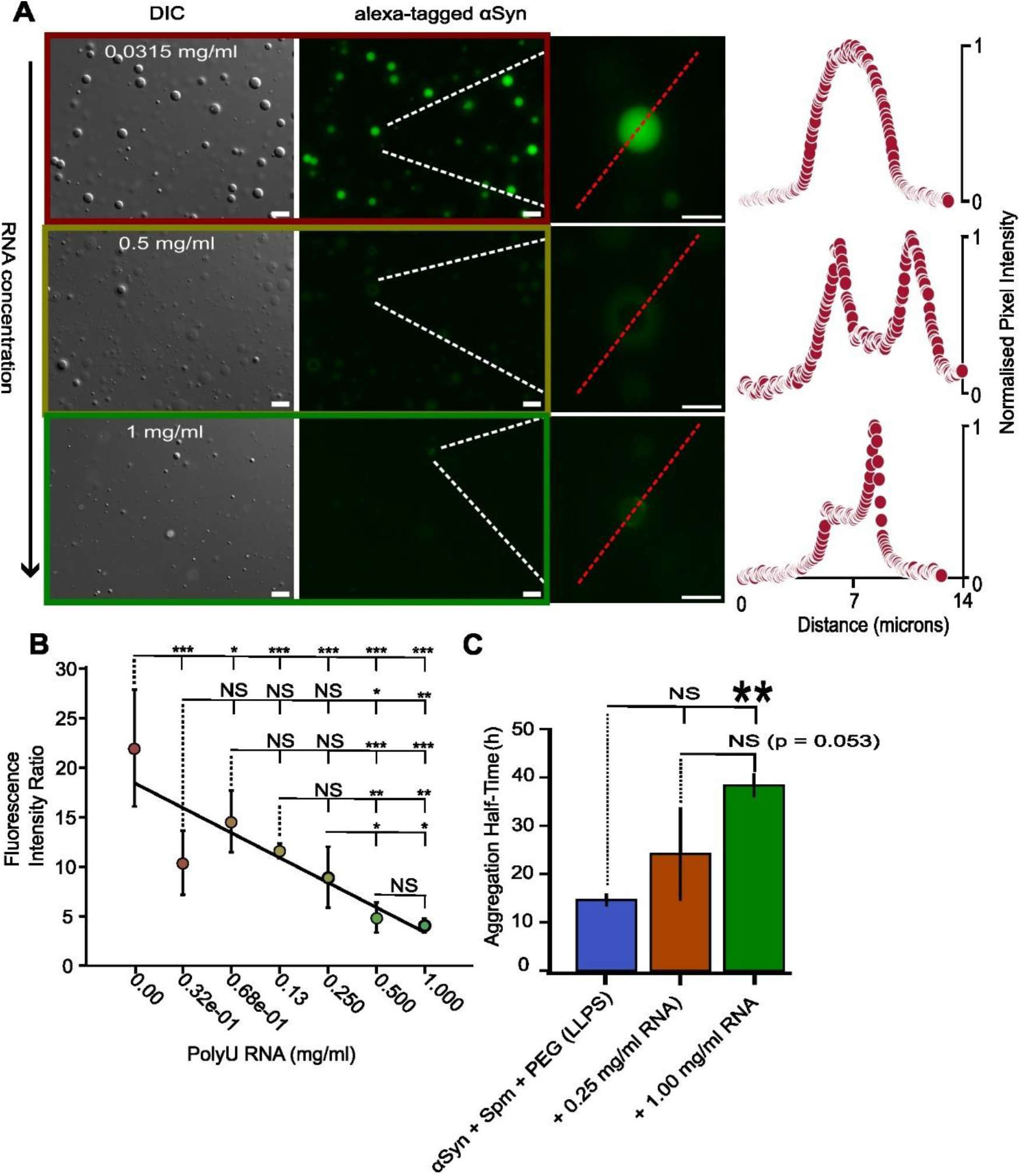
RNA: spermine competitive interactions resolubilises aSyn. **(A)** DIC and fluorescence micrographs of aSyn and spermine with variable polyU RNA concentrations. **(B)** Fluorescence intensity ratio defined as the height of the fluorescence peak within the droplets divided by the averaged background in the vicinity of the droplet. Error bars represent standard deviation and asterisk represent p-value obtained from one-way ANOVA test. **(C)** Aggregation halftime in hours defined as the mean of the inflection points of each replicate fitted with a sigmoid function. Error bars represent standard deviation of the three replicates. Scale bars 5 µm,

RNA-induced weakening of spermine’s interaction with aSyn’s C-terminus and the protein’s subsequent depletion from droplets suggests that RNA might decrease aSyn’s aggregation kinetics in the presence of spermine. Consistent with this hypothesis we found that RNA slows aSyn’s aggregation kinetics in the presence of spermine in otherwise identical solution conditions (**Figure S7C**). With increasing concentrations of RNA, the lag time of aSyn’s aggregation increases substantially relative to the reference phases separation condition (**Figure 7C**).

## Discussion

Abundant cellular molecules are likely to promote, negate and control the formation and regulation of biomolecular condensates in the complex environment of cells ^37^. Here we show that the cellularly abundant natural polyamines, which may exceed millimolar concentrations inside cells ^38^, promote the phase separation of MLO-associated proteins containing acidic domains (**Figures 1-5**).

G3BP1 is a scaffold protein of stress granules which has its saturation concentration for phase separation lowered in the presence of RNA, an interaction that occurs predominantly at the basic termini of the protein ^33^. It was found that removal of the central, long acidic segment of G3BP1 (IDR1) in these conditions enhanced the phase separation of G3BP1 in the presence of RNA. This led the authors to suggest that the acidic disordered region represents an autoinhibitory domain of G3BP1’s phase separation ^19^. However, this is a context-dependent classification since intramolecular interactions which compete with intermolecular interactions (whether RNA or polyamines) that are necessary for the phase separation of the protein will likely increase the saturation concentration. Our results show that the natural polyamine spermine promotes the phase separation of G3BP1 (**Figure 1**).

We further show that spermine increases the clustering and phase separation of aSyn (**Figures 2-3**). aSyn has recently been suggested as a modulator of processing bodies – a type of MLO – and mRNA stability ^39^. Since polyamines are strongly correlated to cellular mRNA there may be a physiological connection between the polyamines and aSyn in processing bodies. However, the effect of the polyamines is expected to be more general since many proteins bear a negative net charge at physiological pH. The results highlight that heterotypic electrostatic interactions even with proteins that are amphiphilic and contain many non-charged residues can still promote phase separation.

Our results suggest that spermine-mediated phase separation accelerates aSyn’s aggregation kinetics (**Figure 4**). This may be especially important considering the increasingly appreciated association of polyamines with Parkinson’s disease ^15^. We also show that the polyamine-promoted phase separation and aggregation depends on RNA concentration, with increasing RNA concentration reducing the attractive interactions between spermine and aSyn and resolubilising αSyn (**Fig. 6-7**). Consistent with a regulatory role of polyamines in the life cycle of biomolecular condensates, polyamines are known to regulate the association of RNA-binding proteins with mRNA via competitive interactions ^40^. These competitive interactions reflect a mechanism that cells may exert to control composition, assembly and disassembly of MLOs.

Acidic domains of proteins which interact with polyamines might be considered analogous to RNA-binding domains of proteins in that they control attraction and repulsion of proteins via electrostatic complementarity. Polyamines, however, are much shorter than most RNA molecules in the cell and are thus less likely to act as a scaffold for phase separation. On the other hand, owing to their multivalency and large size relative to other cellular cations, polyamines have been demonstrated to bridge like-charged ions and nucleic acids together into network-like structures ^41–43^ . Polyamines might therefore be predominantly involved with initial protein and RNA clustering processes as well as in competitive interactions that control assembly and disassembly of MLOs ^44^.

The physiological function of cellular polyamines is intimately tied to the regulation of RNA and stress responses ^10,22,40,45,46^. Polyamines regulate the assembly and disassembly of stress granules in a concentration-dependent manner, and depletion of polyamines enhances stress granule formation in epithelial cells ^36^. However, owing to their small molecule nature, it is challenging to separate direct and indirect effects of polyamines on biomolecular condensates. Since it has been suggested that most of the intracellular polyamine pool is in the bound state with RNA ^6^, and polyamines are directly implicated in cellular stress responses ^22^, it is tempting to suggest that polyamines have direct involvement with stress granules and other RNA-bodies. This involvement might be related to a recently suggested model of MLOs whereby stoichiometry-dependent competition between stickers of macromolecules controls the assembly, disassembly and composition of MLOs ^44^. Our data further suggest that this process can be regulated by acetylation and metabolic conversion of spermine (**Figure 5**), processes that naturally occurs in cells ^30^. Taken together our data suggest a critical role of cellular polyamines in the regulation of biomolecular condensation.

## Methods

### Protein production

aSyn was recombinantly produced in BL21(DE3) Escherichia coli cells (Novagen) as previously described ^47^. Fulllength G3BP1 was obtained from Novus Biologicals (NBP1-50925-50UG). Upon arrival, the sample was dialysed into 50 mM Tris, pH 7.4, 1 mM DTT at 4 °C. The protein was subsequently frozen at -20 °C. G3BP1-peptide was obtained as a powder and was subsequently dissolved in 50 mM Tris buffer, pH 7.4. The sample was then frozen at -20 °C until needed.

### Dynamic light scattering

DLS measurements were conducted at variable temperatures using a DynaPro NanoStar instrument (Wyatt Technologies) and NanoStar disposable microcuvettes. The samples were illuminated with a 120 mW air-launched laser at a wavelength of 662 nm and the intensity of light scattered at an angle of 90° was detected with an actively quenched, solid-state single-photon counting module. Data were acquired with an acquisition time of 5 s with a total of 10 acquisitions per measurement. The hydrodynamic radii were determined using the Dynamics (version 7.10.0.23) software package.

### Turbidity (spectrophotometry)

Turbidity measurements were conducted using a Eppendorf Biospectrometer® kinetic (Eppendorf®) instrument in a 10mm path-length microvolume Quartz Cuvette. The samples were illuminated at a wavelength of 600 nm. Data were acquired in triplicate unless stated otherwise. In conditions of 20% w/v PEG-8000, PEG alone would firstly be measured repeatedly to ensure a constant signal before baselining for future measurements. In between measurements the cuvette was rinsed several times with distilled H^2^O and dried using a steady flow of nitrogen gas.

### Light microscopy

Phase separation experiments were monitored by differential inference contrast microscopy and fluorescence microscopy and were conducted at room temperature unless stated otherwise. Samples were fluorescently labelled on lysine residues using Alexa Fluor 488 Microscale Protein Labelling Kit (Thermo Fisher Scientific, #A30006) according to the manufacturer’s instructions. In all experiments, small amounts of labelled protein, which was insufficient to phase separate by itself, was added to unlabelled protein to the final concentration indicated in the text. If not stated otherwise, the RNA analog polyuridylic acid was used (polyU; Sigma #P9528).

A total of 5 µL of sample was loaded onto a glass slide and covered with a 18 mm coverslip. DIC micrographs as well as fluorescent micrographs were acquired on a Leica DM6B microscope with a x63 objective (water immersion) and processed using ImageJ.

### NMR spectroscopy

For G3BP1-peptide, NMR spectra were recorded at 15 °C on a 800 MHz spectrometer equipped with a triple-resonance cryogenic probe. The protein was dissolved in 50 mM Tris buffer, pH 7.4, with or without 1250 µM spermine, along with DSS for proton chemical shift referencing and 10% D2O. One-dimensional ^1^H NMR and 2D ^1^H-^1^H TOCSY and NOESY spectra were acquired. In addition, ^1^H-^15^N heteronuclear single quantum coherence (HSQC) spectra were recorded. Samples were prepared by dissolving the peptide powder in the NMR buffer before temperature equilibration for 15 minutes inside the spectrometer.

For aSyn, spermine and polyU RNA interaction studies, 600 MHz, 700 MHz and 800 MHz NMR spectrometers (Bruker) equipped with a triple-resonance cryogenic probe were used to record one-dimensional and two-dimensional spectra. Specific details are provided in relevant figure legends, but most often measurements were conducted at 15°C in hepes buffer, pH 7.4, with DSS ^1^H chemical shift referencing and 10% D2O. All data were processed using Topspin 4.1.4 (Bruker) and analysed with CCPNMR 3.1. For calculating the dissociation constant between aSyn and spermine a weighting coefficient of 0.14 for 15N was used.

For the slice-selection experiments of the aSyn: spermine and PEG-8000 phase separated solution as well as the reference samples NMR spectra were recorded at 15°C on a 700 MHz spectrometer equipped with a triple-resonance cyrogenic probe. The 1D spin-echo imaging pulse sequence – or z-profile – was acquired with 2 transients 1 dummy scan and a relaxation delay of 30s. For the spatially selective experiments a WET-DPFGSE pulse sequence was used as previously described ^28^. For this experiment 256 transients and 8 dummy scans were used with a 1.5s relaxation delay.

The conditions for the phase separated sample were 600 µM aSyn, 6 mM spermine, 10% w/v PEG-8000 and 1mM DSS. The sample was transferred to a 3.0 mm NMR Shigemi tube and assembled in a coaxial manner inside a 5.0 mm NMR tube filled with D20 for spectrometer locking. To form a condensate, the coaxial NMR tube was centrifuged at ∼1500 g for many hours at room temperature. Three other samples were measured for the quantification of concentrations in the phase separated system: sample 1 contained 10% w/v PEG-8000 in 50 mM Hepes solution, sample 2 contained 600µM aSyn solution in 50 mM Hepes solution, and sample 3 contained 6 mM spermine in 50 mM Hepes solution.

### Aggregation assays

Aggregation of 50 µM aSyn was performed in 50 mM Hepes, 10mM NaCl and 20% w/v PEG-8000 with or without 500 µM Spermine, pH 7.4. ThT was added to the solution at a final concentration of 50 µM to monitor aggregation kinetics. A total of 100 µL of 50 µM aSyn protein with 50 µM ThT, 20% w/v PEG-8000 and 500 µM spermine was pippeted in a well of 96-well plate (Greiner Bio-one, microplate, 96 well, PS, F-bottom, Chimney well, uClear, black, non-binding, item no-655906). The aggregation assay was performed at 37 °C in a Tecan spark plate reader with double orbital shaking (shaking duration – 1min, shaking amplitude – 6mm, shaking frequency – 54rpm) at an interval of 10 min. An excitation filter at a wavelength of 430 nm with an excitation bandwidth of 35 nm was used to excite ThT. The emission wavelength was set to 485 nm with a bandwidth of 20 nm (manual gain – 40, number of flashes – 30, integration time – 40 us). The Z-position was calibrated using an empty well before starting each experiment. The ThT fluorescence were called using Tecan Spark control software (v 2.2). The analysis of the aggregation data was performed using MATLAB.

Spermine-induced phase separation of aSyn in the presence of either 0.25 or 1.0 mg/ml polyU RNA was conducted in the same manner as above.

### Tranmission electron microscopy

Samples were adsorbed onto 400-mesh carbon-coated copper grids and the buffer removed using filter paper. Subsequently, samples were stained by the addition of 1% uranyl acetate solution and dried with filter paper. The grids were imaged using a Talos L120C G2 electron microscope.

### Statistical analysis

P > 0.05 was considered non-significant. * P <0.05, ** P < 0.01, *** < 0.001 by one-way ANOVA. Statistical analysis was performed in MATLAB (The Mathworks Inc. 2023).

## Author Contributions

M.P. and C.F.P performed NMR spectroscopy. M.P. biochemical experiments, fluorescence microscopy, and phase separation experiments. M-S.C-O. prepared recombinant aSyn. S.B. supervised preparation of fluorescently-labelled aSyn. M.P. and M.Z. wrote the original manuscript. M.P, C.F.P and M.Z. reviewed and edited the manuscript. M.Z. designed the project.

## Funding Sources

M.Z. was supported by the European Research Council (ERC) under the EU Horizon 2020 research and innovation programme (grant agreement No. 787679) and the VolkswagenStiftung (Project-ID AZ 98188).

## Notes

The authors declare no competing interests.

## Acknowledgements

We thank Kerstin Overkamp for solid-phase synthesis of the G3BP1 peptide, and Karin Giller for preparation of fluorescently-labelled aSyn.

## Notes

### Competing Interest Statement

The authors have declared no competing interest.

### Summary of Updates

Arrangement of figures updated and minor errors modified

