## Supplementary Files for "Polyamines promote disordered protein phase separation"

**Figure S1**

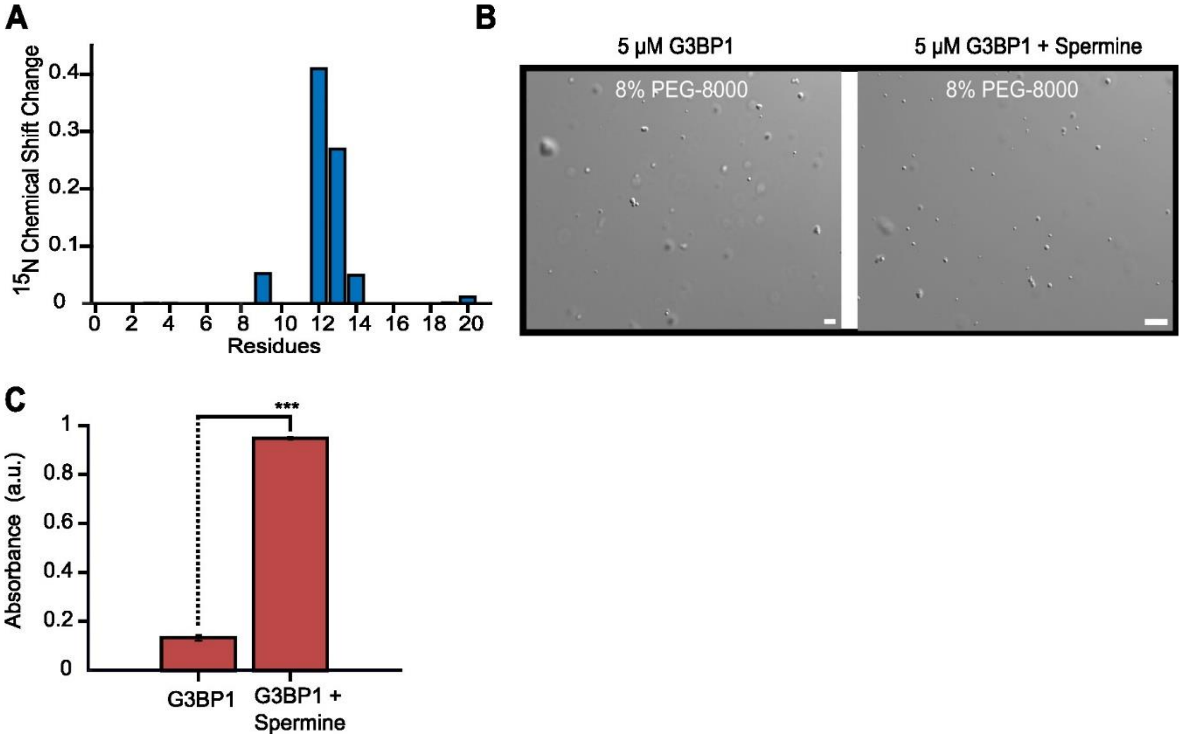


**Spermine interacts with G3BP1-pep and phase separates full-length G3BP1.** 15N chemical shift perturbations of G3BP1-pep in the presence of spermine **(A)** and spermine-induced phase separation with full-length protein **(B)** and turbidity **(C).**

**Figure S2**

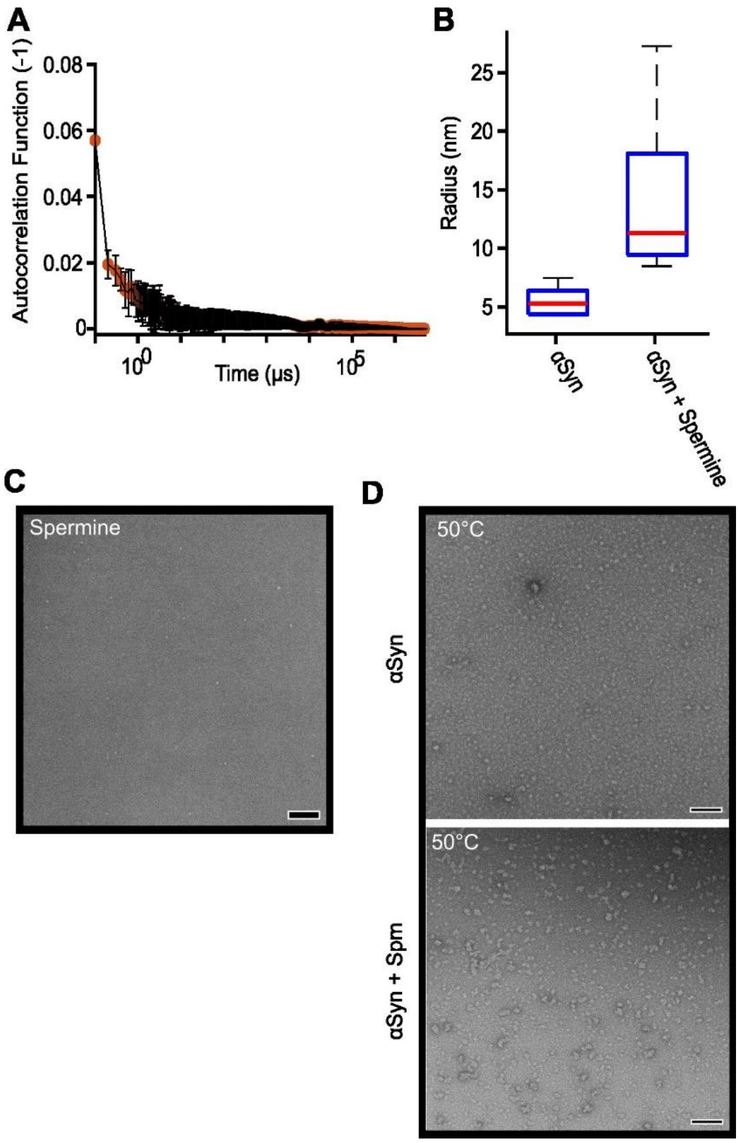


**Spermine and aSyn form clusters. (A)** Auto-correlation function of spermine. **(B)** Mass-weighted average radii of αSyn and αSyn + Spermine. Bottom and top edges of the box represent $25^{th}$ and 75th percentiles, respectively. The whiskers extend to the most extreme data points which are not outliers. Red bar represents the median of the samples. **(C)** TEM micrograph of spermine in Tris buffer. **(D)** 250 µM αSyn and αSyn + 2500 µM Spermine micrographs recorded after a 30m incubation at 50°C.

**Figure S3**

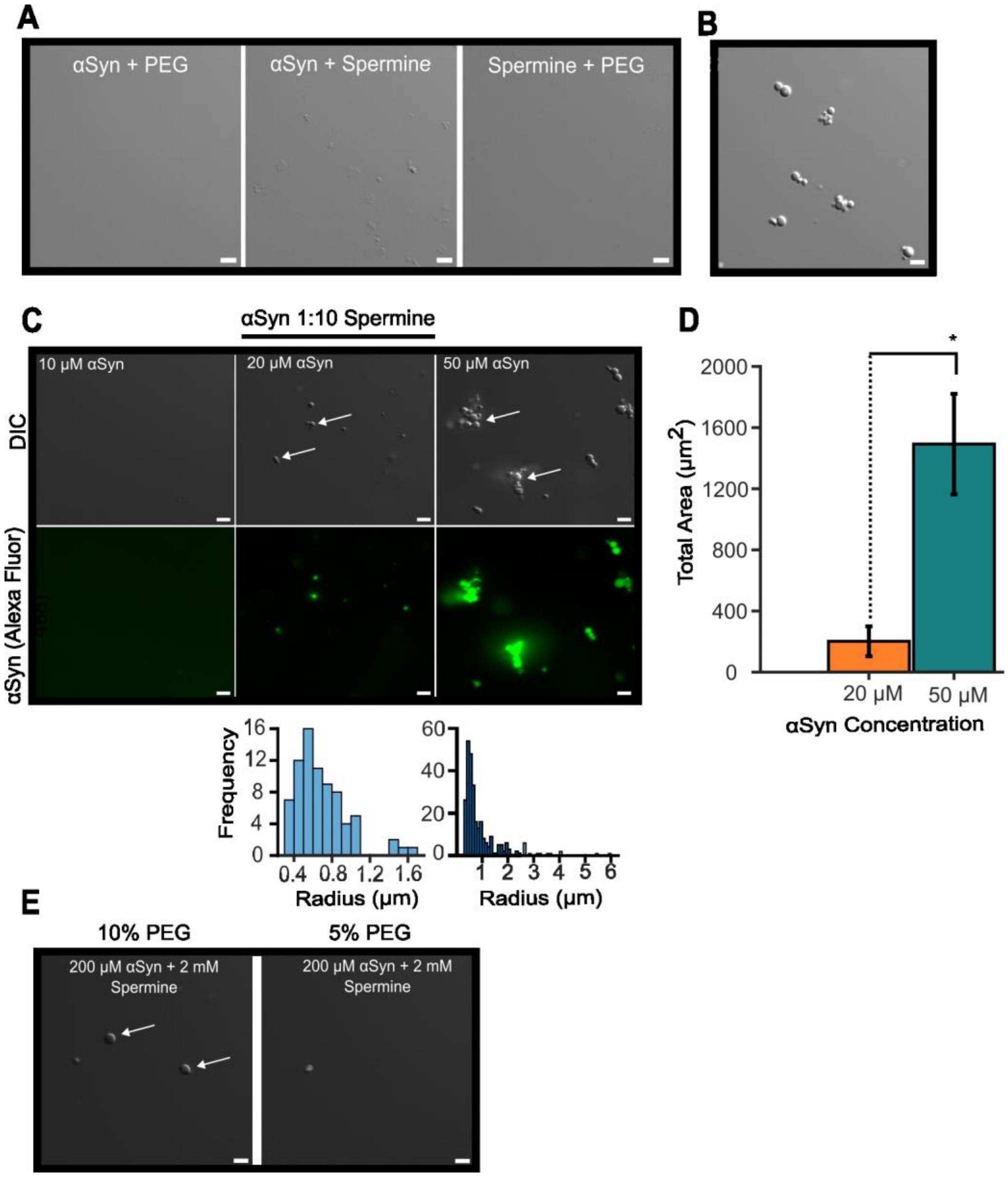


**DIC and fluorescent microscopy images of αSyn, spermine and PEG at various stoichiometries. (A)** 50 µM αSyn mixed with either (left) 20% w/v PEG-8000, or (middle) 500 µM spermine. (right) 500 µM spermine + 20% w/v PEG-8000. Scale bar is 5 µm. **(B)** 50 µM αSyn with 500uM spermine and 20% w/v PEG-8000 without fluorescent labelling. Scale bar is 5 µm. **(C)** αSyn + 1:10 molar ratio of spermine at different initial αSyn concentrations with radius histograms below the images. **(D)** Total area of fluorescent droplets. **(E)** αSyn and spermine mixtures at different PEG concentrations.

**Figure S4**

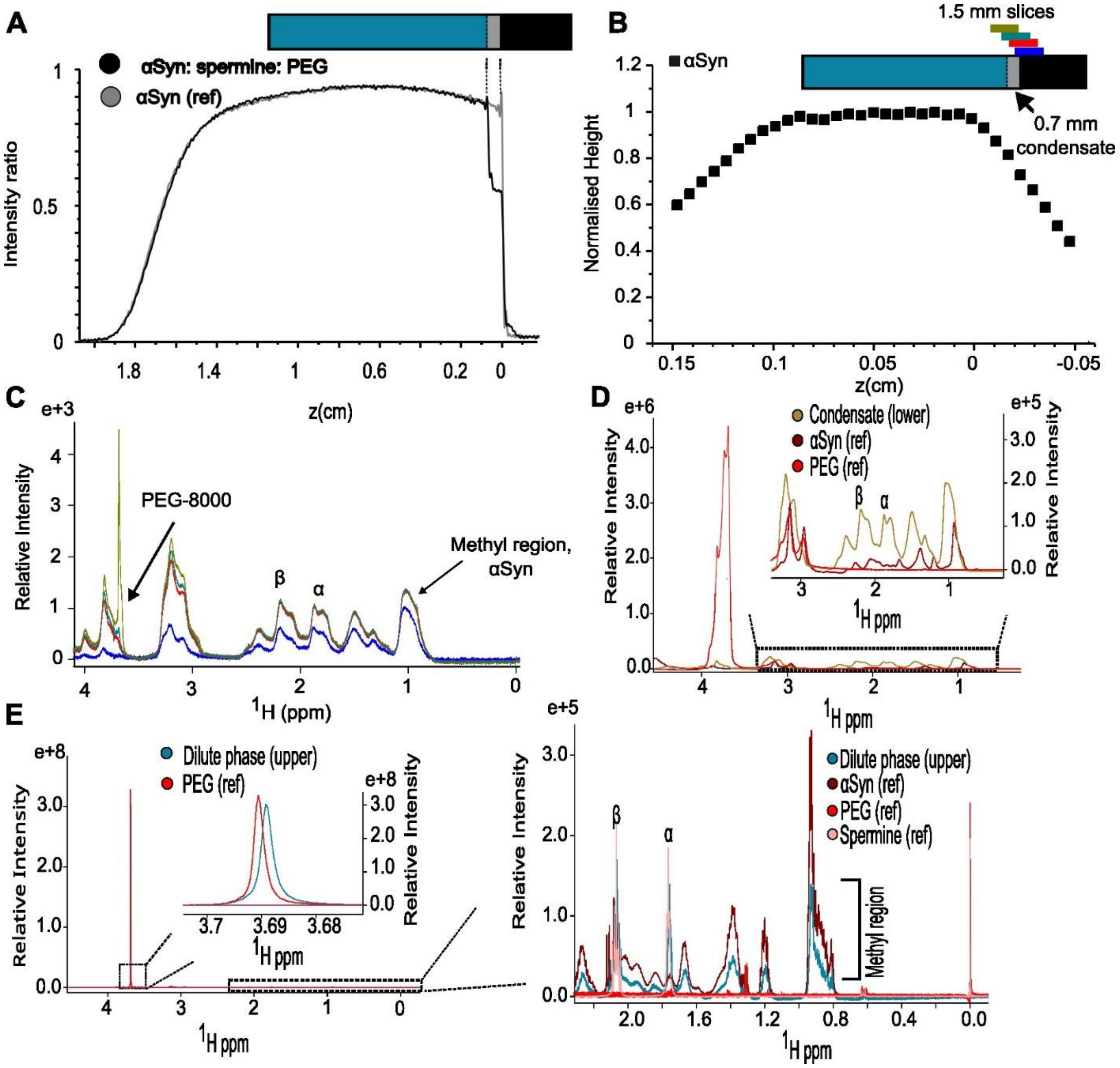


**Reference NMR spectra for quantification. (A)** Spin density of water along Z-axis of NMR tube. **(B)** Intensity of methyl region at different positions (offset frequencies) along the Z-axis of the NMR tube. Each data point corresponds to the methyl intensity from a single slice. **(C)** 1H NMR spectra at different Z-positions (offset frequencies). Reference 1H NMR spectra at the positions (offset frequencies) corresponding to either the condensate **(D)** or the dilute, upper phase **(E).**

**Figure S5**

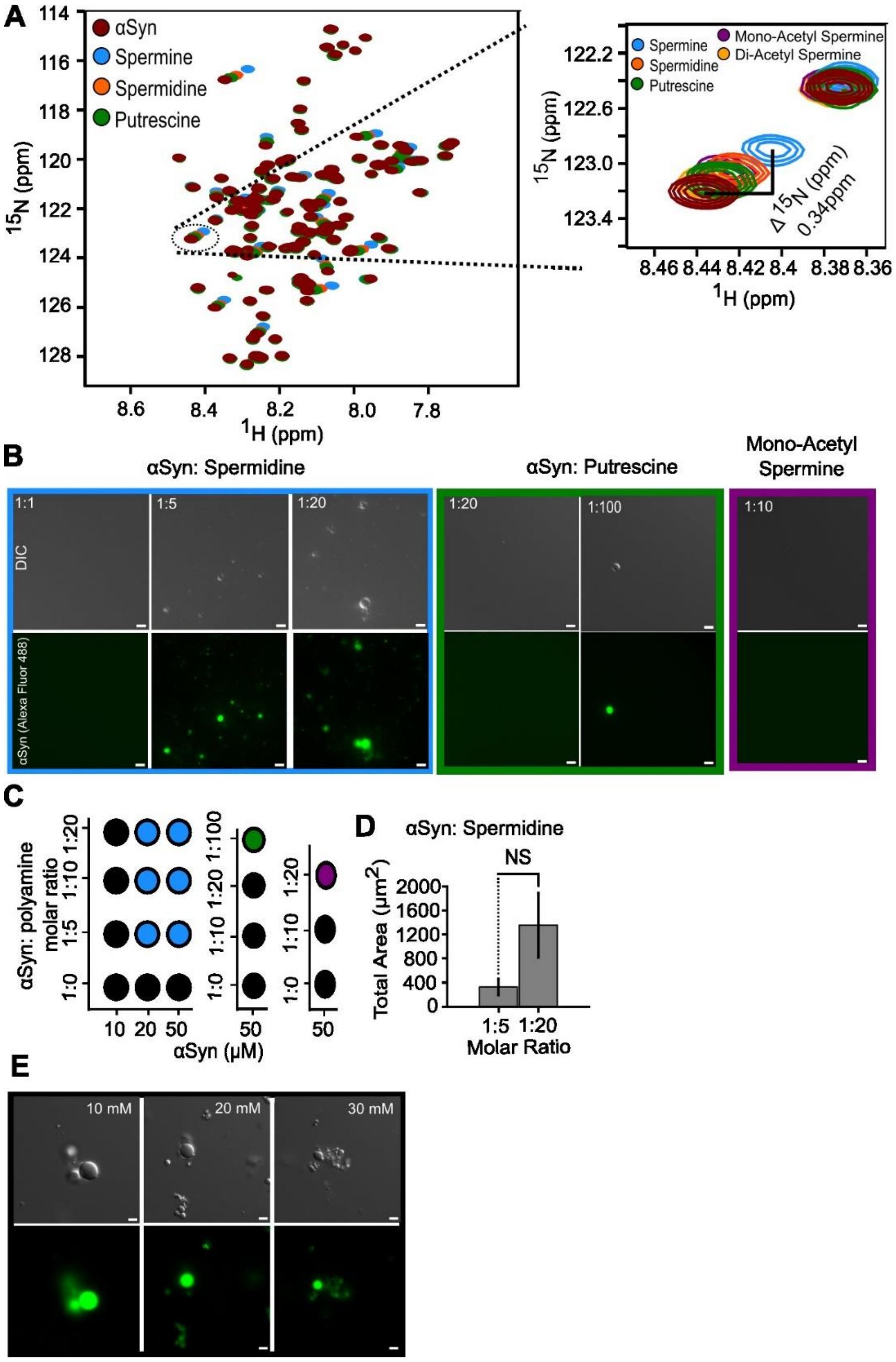


**Microscopy images of the concentration dependence of polyamine-dependent phase separation.** **(A)** 1H-15N HSQC of aSyn in the presence of different polyamines. **(B)** Microscopy images of aSyn in the presence of different concentrations of various polyamines. **(C)** Stoichiometry diagrams of polyamine and aSyn phase separation. **(D)** Total fluorescent area of spermidine and aSyn phase separation. **(E)** Microscopy images of aSyn and spermine phase separation with different NaCl concentration.

**Figure S6**

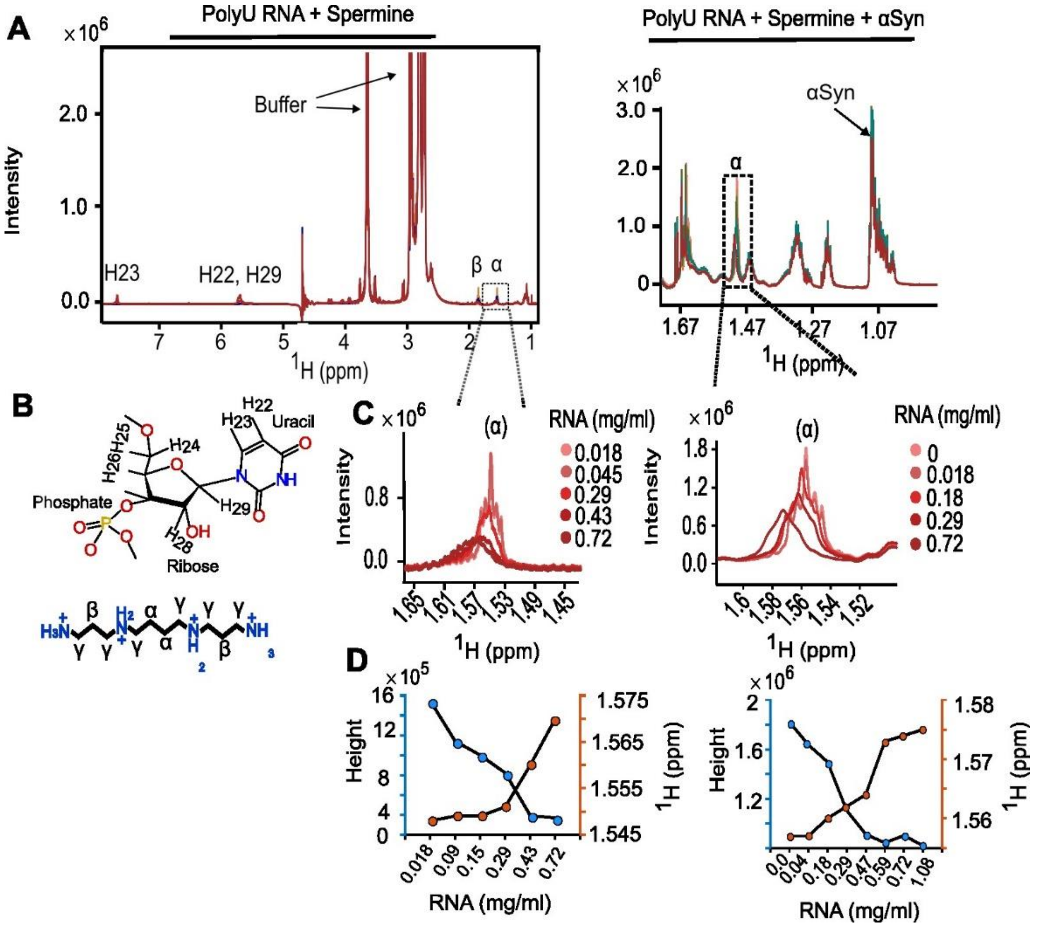


**1H NMR spectra of RNA-induced changes to spermine's chemical shift and lineshape with or without aSyn present. (A)** 1H NMR spectra of PolyU RNA and spermine (left) and PolyU RNA, spermine and aSyn (right). **(B)** chemical structures of a monomer of polyU RNA and spermine with relevant assignments. **(C)** inset on spermine's chemical shift changes with increasing amounts of polyU RNA without (left) or with (right) aSyn present in solution. **(D)** Spermine's α chemical shift and height changes.

**Figure S7**

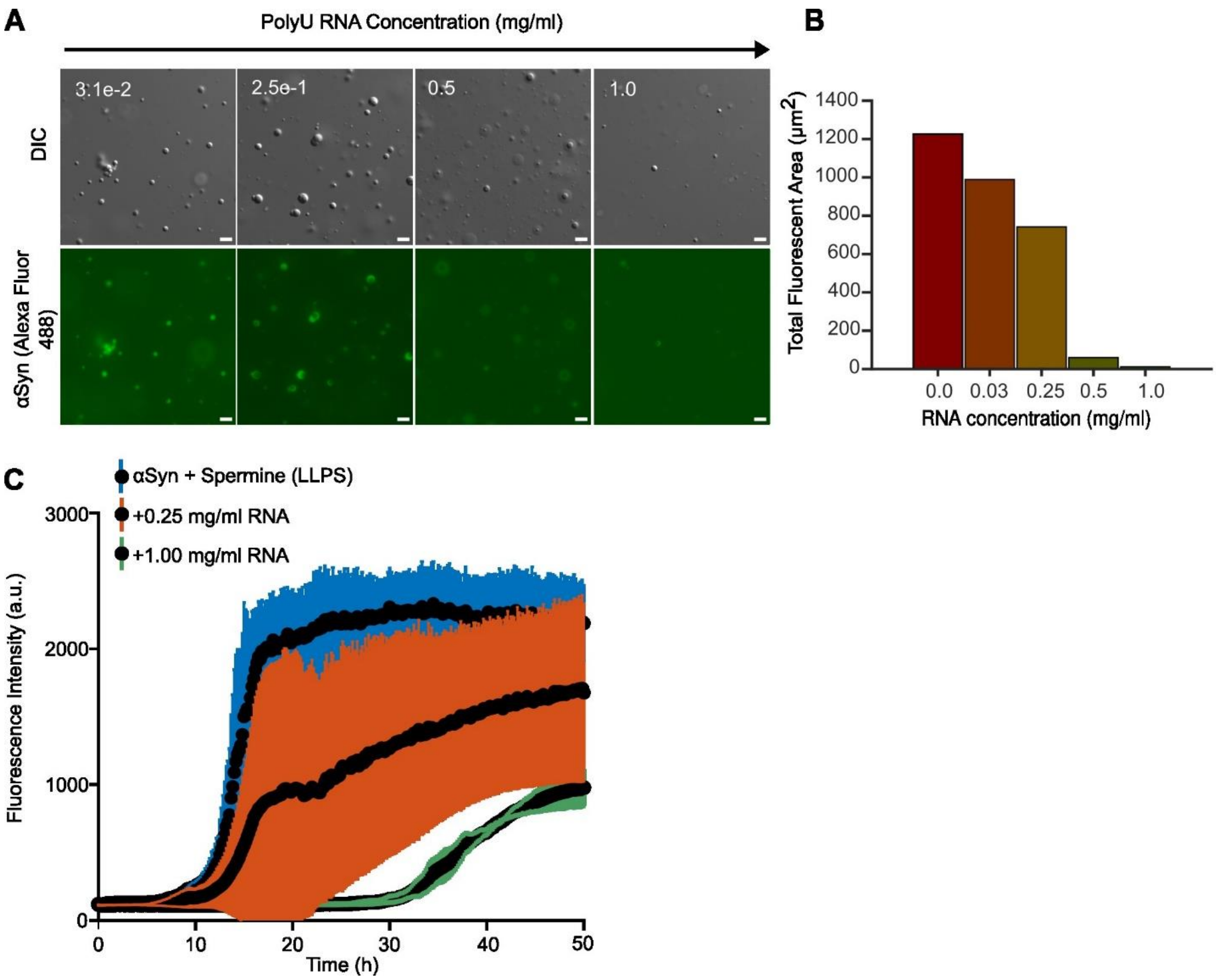


**RNA resolubilises aSyn and reduces its aggregation rates in phase separation conditions. (A)** Microscopy images of (labelled) aSyn and spermine at a molar ratio of 1: 10 with increasing RNA concentration and corresponding fluorescent areas **(B). (C)** Th-T curves of aSyn in phase separation conditions (blue) and with increasing amounts of RNA (orange, green).
